# More extinction driven by Red Queen in smaller habitats

**DOI:** 10.1101/2024.05.22.595421

**Authors:** Xiao Liu, Quan-Guo Zhang

## Abstract

Populations in antagonistic coevolutionary interactions may “run or die”, and their fates are determined by their evolutionary potential. The asymmetry of evolutionary speed between coevolving partners, e.g., resulting from genetic constraints, can be mitigated in larger populations. We hypothesize more frequent extinction driven by antagonistic coevolution when habitat size declines. In bacterium-virus systems, viruses (the consumers) typically suffer an evolutionary disadvantage due to constraints of genetic variation; and this pattern may apply to host-parasite interactions in general. Here in our experiment with the bacterium *Pseudomonas fluorescens* SBW25 and its lytic phage virus SBW25Φ2, the likelihood of viral extinction was greater in smaller habitats. Among viral populations that did persist, those from small habitats showed lower infectivity and their coevolving bacterial populations had greater densities. Therefore, the impact of habitat size reduction on biodiversity could be exacerbated by coevolutionary processes. Our results also lead to a number of suggestions for biocontrol practices, particularly for evolutionary training of phages.

## INTRODUCTION

Organisms are usually engaged in antagonistic species interactions; evolving enemies and competitors become continuously deteriorating biotic environments [1-3]. Red Queen dynamics characterized by perpetual evolutionary changes are believed to be very common, which may also end up with extinction of populations that “lose” the game, or evolutionary stasis where interacting populations show no further evolutionary changes or extinction and speciation [4-9]. Analysis of extinction dynamics in fossil records has suggested antagonistic coevolution as a major reason for taxon extinction, though its contribution relative to physical environmental changes has been controversial [2, 10-14]. The role of antagonistic evolution for driving population extinction has also been demonstrated in a number of experimental studies with model systems [15-18].

The asymmetry in the evolutionary potential between coevolving partners could be mitigated under certain ecological conditions but become apparent when environment changes [19-24]. Take bacterium-virus interactions as an example, viral evolution is usually constrained by mutational supply, presumably because of the intrinsic difference in the evolvability between bacterial resistance and viral infectivity mechanisms. The former can be achieved by a number of changes including alteration or loss of viral receptors while the latter requires much more specific changes that may need stepwise acquisition of multiple mutations [25-28]. Viruses also have smaller genomes and thus lower availability of mutations for changing infectivity profiles; and their burst mode of replication is associated with population bottlenecks which reduce effective population sizes. Consistent with this notion, gene flow introducing genetic variation into coevolving populations was found to enhance viral, relative to bacterial, evolution [29-31]. Meanwhile, bacterial evolution often depends on the fitness costs associated with resistance mechanisms such as decreased growth rate [20, 32]. Environmental factors that alleviate such fitness costs, e.g. increased nutrient availability, may accelerate bacterial resistance evolution [33-36] and even drive coevolving viral populations extinct [37, 38].

Here we consider the consequences of habitat size for the outcome of antagonistic coevolution. Habitat size reduction are now causing loss of biodiversity and ecosystem functioning [39, 40]. We hypothesize that smaller habitat size can increases the chance for antagonistic coevolutionary interactions to manifest as asymmetrical and end up with population extinction. The asymmetry in evolutionary potential between coevolving partners may be mitigated in large habitats, as increasing mutational supply has diminished benefit for evolutionary adaptation when population sizes exceed certain thresholds [41, 42]. For bacterium-virus systems where the viral evolution is more limited by mutational constraints, increased habitat sizes are more effective for enhancing viral evolutionary potential but has limited effect on bacterial evolution. We therefore predict that viral extinction driven by bacterial resistance evolution be more likely in smaller habitats.

We carried out an experiment with a model bacterium-virus system, *Pseudomonas fluorescens* SBW25 and its lytic phage SBW25Φ2. This pair of species may show arms race-like or fluctuating selection dynamics of coevolution, depending on the time course of coevolution and environmental factors [23, 43, 44]. Phage extinction driven by bacterial resistance evolution in this system has been documented in a number of studies [16, 17, 37, 38, 45], while it is also clear that the phage can always maintain a viable population when continuously evolving with susceptible bacteria supplemented by investigators [46-49]. The present study examined phage extinction dynamics, evolution of infectivity ranges and impacts on bacterial population densities in habitats of different sizes, and found that smaller habitat size exacerbated the disadvantage of phages in coevolution.

## MATERIAL AND METHODS

### The bacterium-virus system and culture conditions

The present study used the bacterium *Pseudomonas fluorescens* SBW25 [50], and its lytic bacteriophage virus SBW25Φ2 [51]. Cultures were grown at 26 °C in static microcosms, centrifuge tubes with lossened lids that contained certain volumes of LB broth (10 g L^-1^ tryptone, 5 g L^-1^ yeast extract, and 10 g L^-1^ NaCl). LB agar plates were used for bacterial density measurement and phage infectivity assays. Culture samples were stored at -80°C in 40% (vol/vol) glycerol.

### The coevolution experiment

Three types of centrifuge tubes were used to set up microcosms of different habitat sizes (Fig. S1; purchased from Hengchao Limited, Nantong, China). They were all U-bottomed; and the inner diameters were 12, 17 and 37 mm, corresponding to top surface area of 113, 227 and 1075 mm^2^, respectively. Cultures were grown in those centrifuge tubes loaded with liquid media of 20 mm depth (distance from the center of bottom to upper surface). The volumes of nutrient media in the three types of tubes were approximately 1.25, 2.20 and 10.00 mL, respectively. We used habitats of the same depth of liquid broth, with different top surface areas, to rule out the possibility that oxygen limitation confounded with the habitat size effect. Hereafter habitat size was referred to as their top surface area.

Twelve replicate evolution lines were established for each of the three habitats. Every microcosm was initially inoculated with approximately 10^7^ mL^-1^ of bacterial cells and 10^4^ mL^-1^ of phage particles. Cultures were then propagated for 16 transfers, one transfer every two days. At each transfer, 1% of cultures were added to fresh media. We measured bacterial densities at transfer 1 and 16, by spreading culture dilutions on LB agar plates and counting colony forming units (CFUs) after two days of incubation.

### Inference of phage persistence

Phages were isolated from each culture at transfer 1 and 16. Specifically, a mixture of 0.4 mL of culture and 0.04 mL of chloroform was vortexed to lyse bacterial cells, and centrifuged at 15,000 g for 2 min to pellet the bacteria debris, leaving a suspension of phage in the supernatant. We measured plaque forming units (PFUs) on lawn of the ancestral bacterial strain. Dilutions of each phage extract were spotted on plates made with a mixture of soft agar media (LB broth with 7 g L^-1^ of agar) and the ancestral bacterial cells, and PFUs were counted after 24 h incubation. The PFUs measurement indicated phage presence in all microcosms at transfer 1, while PFUs were not detectable for a number of phage populations at transfer 16. The failure to form plaques on the lawn of ancestral bacterium may not confirm the extinction of a phage population. During the coevolution with host bacteria, some phage populations may lose the ability to infect the ancestral bacterium but can infect bacterial genotypes emerging during coevolution [34]. For phage extracts from transfer 16, we also measured their infectivity against a number of bacterial isolates (see below “measurement of infectivity ranges”). A phage population was considered as extinct if both its PFUs on lawn of the ancestral bacterium was zero and its infectivity scores were zero.

### Measurement of infectivity ranges

For each phage extract from transfer 16, two infectivity range scores were measured by bacterium-phage challenge assays on agar plates [51]. First, infectivity against a common bacterial pool was measured. A pool of 24 reference bacterial isolates were set up, each isolate from a randomly chosen microcosm. Colonies of those reference bacterial isolates were picked up from agar plates and inoculated into liquid media. We streaked a line of phage extract (20 μL) on an agar plate; after the phage lines dried, we streaked culture of each of reference bacterial isolates across the phage lines. After 24 h of incubation, we checked the growth of the bacterial lines. Inhibition of bacterial growth indicated phage infectivity. Infectivity range of each phage population was defined as the proportion of susceptible bacterial isolates among the total of 24.

Second, infectivity against within-microcosm bacteria was measured for each phage population, by determining the susceptibility of 20 bacterial isolates from the local bacterial population that had coevolved with the specific phage population. For a number of phage populations whose persistence was confirmed, the score of infectivity against within-microcosm bacteria was recorded as zero. Here the zero score was in fact “< 0.05”, as our assays of phage against within-microcosm bacteria were based on 20 bacterial colonies per microcosm. In such cases, phage may have been maintained because of susceptible bacterial genotypes that were rare and undetected by our phage infectivity assays.

### Data analysis

The R environment was used for data analysis [52]. The probability of phage persistence was compared among habitats using Unconditional Exact Test [53]. Infectivity data were analyzed using Wilcoxon tests. For multiple comparisons, *P* values were adjusted using the Benjamini-Hochberg method. The differences in bacterial density (log-transformed) among habitats were analyzed using ANOVA and subsequent Tukey multiple comparisons. The relationship between bacterial density and phage infectivity across microcosms of all habitats was analyzed using a general linear model; and automatic model simplification was carried out using the “step ()” function.

## RESULTS

Phages were detected in all microcosms in the beginning of the experiment (transfer 1). Until the end of the experiment (transfer 16), phages persisted in all the 12 large microcosms (1075 mm^2^). This was not the case in the small microcosms (113 mm^2^) where 8 out of 12 phage populations survived, nor in the medium-size microcosms (227 mm^2^) where 5 out of 12 phage populations survived (Fig. 1a; Exact test, 8/12 versus 12/12, *P* = 0.038; 5/12 versus 12/12, *P* = 0.002; 8/12 versus 5/12, *P* = 0.308; adjusted *P* values for the above three: 0.057, 0.006, and 0.308).

**Figure 1.**
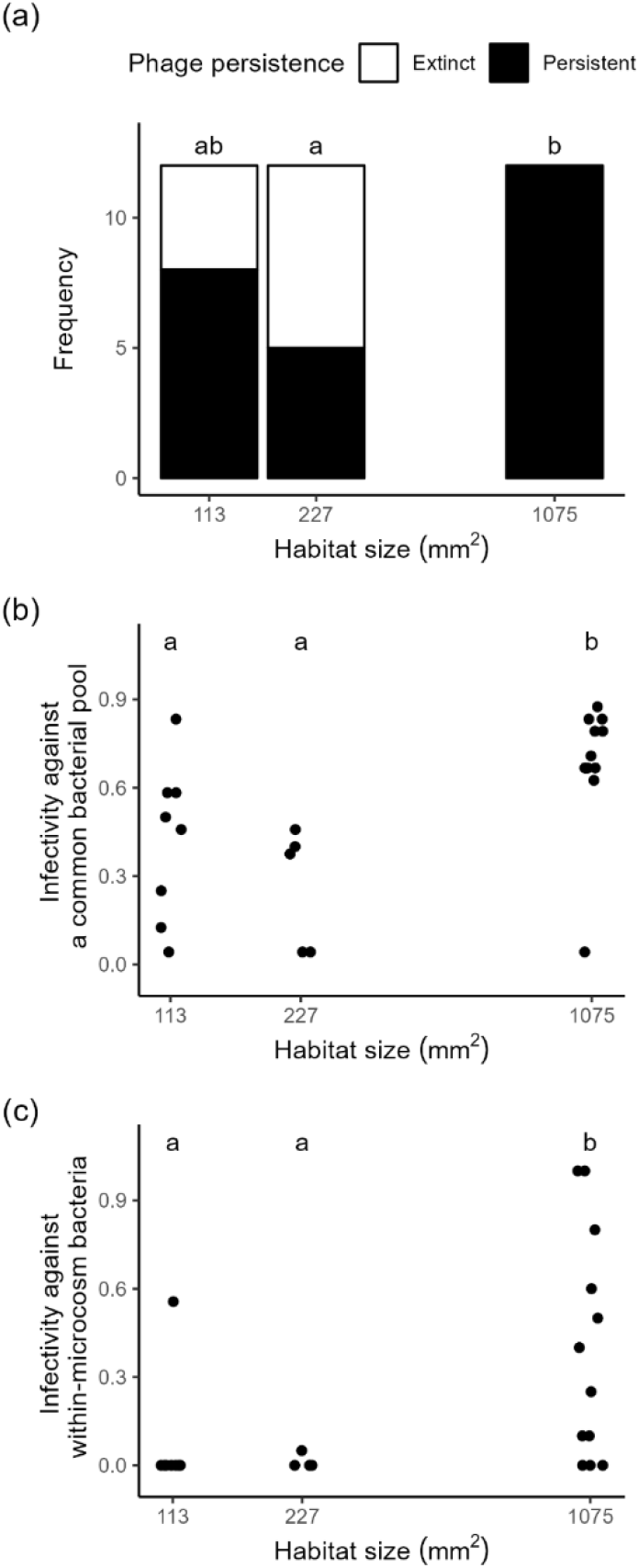
The probability of phage persistence (a) and two infectivity measures of the persisting phage populations (b-c). Scatter points in panels (b-c) were jittered along the horizontal direction to avoid overlapping. In each panel, habitats annotated with a same letter showed no significant difference (*P*_adj_ > 0.05; *P* values adjusted using the Benjamini-Hochberg method). Comparisons in panel (a) were carried out using Exact tests (note that unadjusted *P* values suggested a significant difference between the 113 and 1075 mm^2^ habitats); and those in panel (b-c) were based on Wilcoxon rank sum tests.

For each phage populations that persisted until the end of the experiment, two infectivity scores were measured: infectivity against a common bacterial pool, and infectivity against within-microcosm bacteria. Phages from the large microcosms exhibited greater infectivity against the common bacterial pool than the medium- or small-size microcosms, while no significant difference was found between the latter two types of habitats (Fig. 1b; Wilcoxon rank sum test, small versus large habitats, *W* = 16.5, *P* = 0.016; medium versus large, *W* = 4, *P* = 0.006; smaller versus medium, *W* = 10.5, *P* = 0.184; adjusted *P* values: 0.024, 0.021, and 0.184). A similar pattern was found for the infectivity score measured against within-microcosm bacteria (Fig. 1c; small versus large habitats, *W* = 18.5, *P* = 0.017; medium versus large, *W* = 9, *P* = 0.025; small versus medium, *W* = 21, *P* = 0.907; adjusted *P* values: 0.038, 0.038, and 0.907). Across all the surviving phage populations, infectivity against within-microcosm bacteria was greater than that against the common bacterial pool (the difference between the two scores tested against the expected values of zero, Wilcoxon singed rank test, *V* = 308, *P* < 0.001).

Bacterial density did not show significant differences among the three types of habitats at transfer 1 (Fig. 2a; ANOVA, *F*_2,22_ = 2.076, *P* = 0.141). At the end of the experiment (transfer 16), however, the mean bacterial densities of the small- and medium-size microcosms were 6.74- and 4.40-fold of that of the large-size habitats, respectively (Fig. 2b; *F*_2,22_ = 12.22, *P* < 0.001; Tukey multiple comparisons, small versus large habitats, *P*_adj_ = 0.031; medium versus large, *P*_adj_ = 0.117; small versus medium, *P*_adj_ = 0.966).

**Figure 2.**
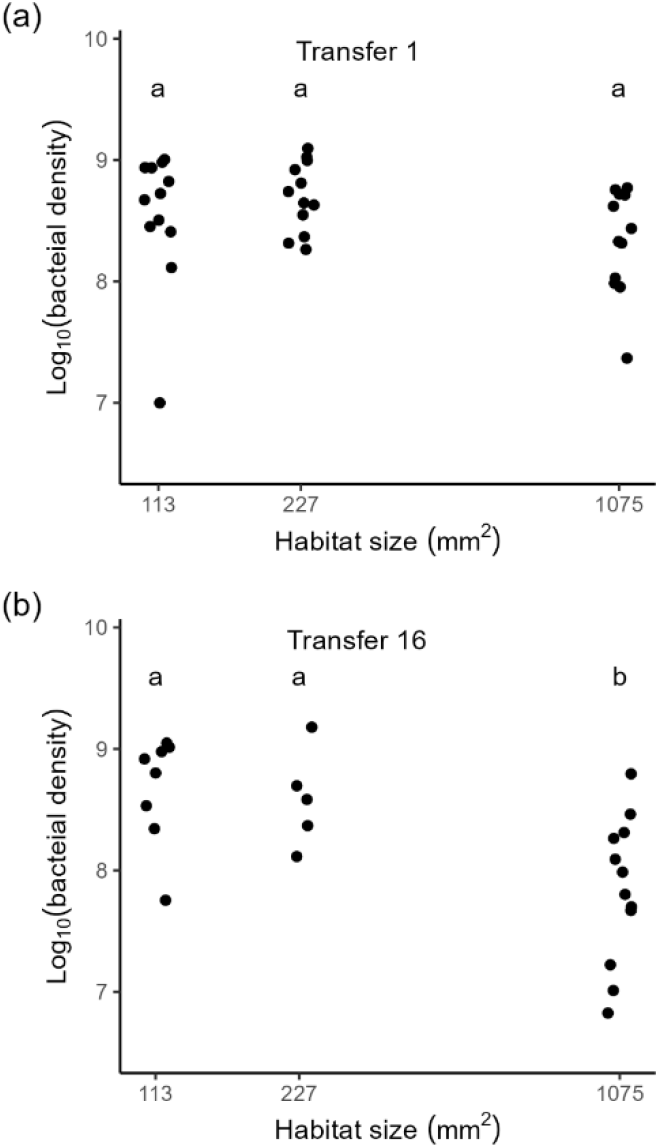
Bacterial density (colony forming units per mL, CFUs) of microcosms at the beginning (a) and the end (b) of the experiment. Here are shown microcosms with phage persistent. In each panel, habitats annotated with a same letter showed no significant difference (Tukey multiple comparisons based on ANOVA, *P*_adj_ > 0.05).

Across all microcosms with phage persistent at transfer 16, bacterial density was negatively correlated with phage infectivity against the common bacterial pool (Pearsons, *r* = -0.451, *df* = 23, *P* = 0.023), or phage infectivity against within-microcosm bacteria (*r* = -0.458, *df* = 23, *P* = 0.021). While the two infectivity measures were positively correlated (Fig. 3; *r* = 0.657, *df* = 23, *P* < 0.001), infectivity against within-microcosm bacteria was a better predictor for bacterial density. For a linear model with the two infectivity scores as explanatory variables for bacteria density (Type II sum of square test, infectivity against the common bacterial pool, *F*_1,21_ =1.17, *P* = 0.292; infectivity against within-microcosm bacteria, *F*_1,21_ = 1.37, *P* = 0.255; interaction, *F*_1,21_ = 1.22, *P* = 0.282), model simplification led to a minimum adequate model with infectivity against within-microcosm bacteria as the only explanatory variable (*F*_1,23_ = 6.12, *P* = 0.021; Fig. 3).

**Figure 3.**
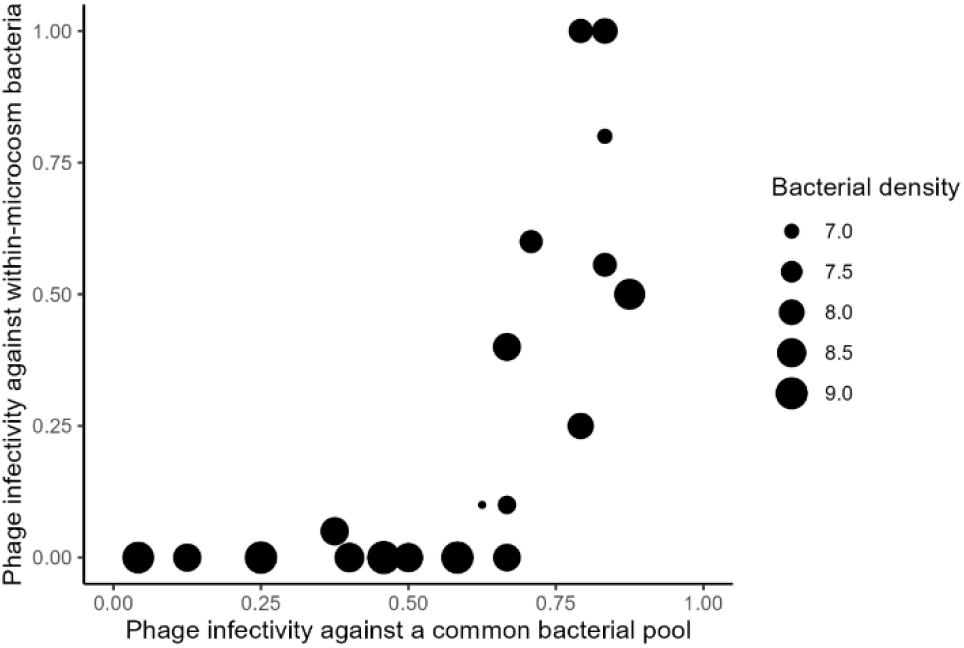
A plot of two infectivity measures of phage populations at the end of the experiment (transfer 16), with the densities of their coevolving bacterial populations (log_10_ (CFUs mL^-1^)) mapped as size of scatter points.

## DISCUSSION

The consequences of larger habitat sizes for maintaining species diversity and ecosystem functioning have long been noticed, and may be attributable to the “more individuals” and “greater heterogeneity” effects [54-58]. Decreases in habitats size caused by human activities are predicted to cause species loss, and the impacts would be more severe for organisms of higher trophic levels [59-61]. As it takes time for extinction events to occur, many species that are now surviving may have already been doomed, and such “extinction debt” will be paid in the not-too-distant future [62, 63]. Here our study suggests a new mechanism for habitat reduction to impair species diversity: exacerbated asymmetry in antagonistic coevolution. Taking this evolutionary mechanism into account would predict more species extinction in destructed habitats.

We formulized a hypothesis for bacterium-virus systems based on two assumptions. First, evolution is more constrained by mutational supply in viruses, relative to their host bacteria. Second, increasing population size in larger habitats has diminished benefit for evolutionary speed, with the asymmetry of evolutionary potential between coevolving partners mitigated. We predict that the evolutionary disadvantage of viruses would be exacerbated in smaller habitats.

In our experiment with a model bacterium-phage system, the evolutionary disadvantage of phages, relative to bacteria, was implied by that score of infectivity against a common bacterial pool was larger than infectivity against within-microcosm bacteria. This pattern of local maladaptation for phages suggests that bacterial populations can better respond evolutionarily to phages, than vice versa, in local habitats. This finding was consistent with earlier studies of our bacterium-phage system [29, 30] and some other host-parasite systems [64]. The evolutionary disadvantage of phages appeared more pronounced in smaller habitats. Our phage populations had a greater probability of extinction when coevolving with bacteria in habitats of smaller sizes (Fig. 1a). Among phage populations that persisted, those from smaller habitats had lower infectivity, and showed weaker controlling effect on host population density (Fig. 1b-c; Fig. 2b). It was also clear that infectivity of local phage populations was a significant predictor of bacterial density (Fig. 3). This was consistent with the expectation that enhanced phage infectivity evolution improve top-down control on bacterial populations, as phages with larger infectivity ranges may directly cause greater bacterial mortality, or drive the evolution of high-level bacterial resistance with larger fitness costs [34, 43].

It is noteworthy that mutational constraint is only one of the possible mechanisms by which habitat size modulate antagonistic coevolution. Decreased population sizes in smaller habitats may also limit the efficiency of selection, relative to drift, particularly for the coevolutionary partners that already have smaller effective population sizes [65, 66]. Smaller habitats may also exhibit lower within-habitat heterogeneity [55, 58] and thus greater extent of population homogenization that may increase the likelihood of asymmetrical coevolution [16, 67].

The consequences of habitat size reduction for asymmetrical coevolution and species extinction observed in our bacterium-phage system may apply to host-parasite interactions in general. Among the two assumptions of our hypothesis, the first is stronger mutational constraints in viral, relative to bacterial, evolution. Such asymmetry may be true for many host-parasite interactions. Though parasites may have large census population sizes, their genetic diversity is often low because of, e.g., extreme population bottlenecks associated with host transmission and asexual reproduction [65, 68]. Prevalent mutational constraints for parasite evolution were implied by a previous meta-analysis that found enhanced parasite local adaptation to hosts when parasites had greater gene flow rates than hosts [69]. The second assumption, diminished benefit of increasing population size for evolution, is expected to hold generally true [41, 42].

Parasitism is arguably the most common consumer strategy [70, 71]. Parasites are critical for ecosystem health, as they contribute to the maintenance of genetic variation in populations and species diversity in communities [72-76]. Extinction of parasites driven by host immunity responses and evolutionary changes in defenses could be frequent, though which may easily be overlooked. Our results suggest more parasite extinction in destructed habitats, which deserves attention from ecologists and conservation biologists. A better understanding of host-parasite coevolution is also helpful for biocontrol practices. Suggestions from our results include to introduce biocontrol agents from larger habitats, and to adopt biocontrol programs on large enough farm lands. Specifically for phage applications, we suggest using as large as possible culture systems in evolutionary training programs that are aimed at obtaining more infective phage materials [47, 77-79].

## Acknowledgements

This study was funded by the National Natural Science Foundation of China (32371687).

## Author contributions

XL and QGZ designed study; XL performed experiments; XL and QGZ analyzed data; all authors wrote the paper.

## Data availability

Data associated with this study are available at figshare (https://figshare.com/s/17ee9b913fadc2680b55).

## Supplementary information

**Figure S1.**
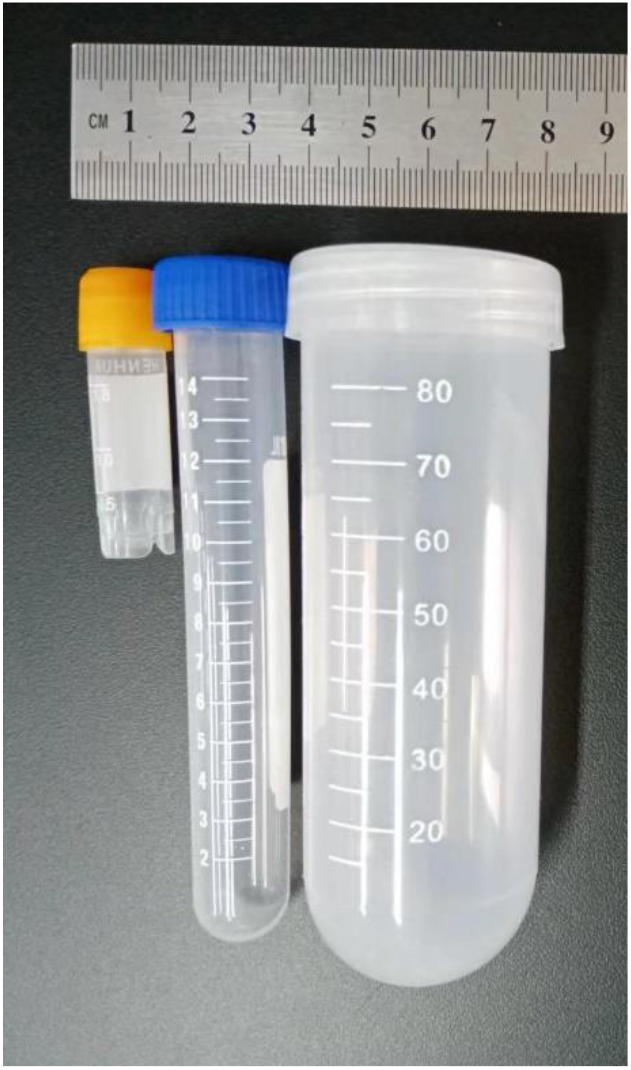
Centrifuge tubes used to set up microcosms. The inner space of those tubes was U-bottomed. The inner diameters of the 2-, 15- and 100-mL tubes were 12, 17 and 37 mm, respectively.

## Notes

### Competing Interest Statement

The authors have declared no competing interest.

